# CONTAIN: An open-source shipping container laboratory optimised for automated COVID-19 diagnostics

**DOI:** 10.1101/2020.05.20.106625

**Authors:** Kenneth T. Walker, Matthew Donora, Anthony Thomas, Alexander James Phillips, Krishma Ramgoolam, Kjara S Pilch, Phil Oberacker, Tomasz Piotr Jurkowski, Rares Marius Gosman, Aubin Fleiss, Alex Perkins, Neil MacKenzie, Mark Zuckerman, Davide Danovi, Helene Steiner, Thomas Meany

## Abstract

The COVID-19 pandemic has challenged diagnostic systems globally. Expanding testing capabilities to conduct population-wide screening for COVID-19 requires innovation in diagnostic services at both the molecular and industrial scale. No report to-date has considered the complexity of laboratory infrastructure in conjunction with the available molecular assays to offer a standardised solution to testing. Here we present CONTAIN. A modular biosafety level 2+ laboratory optimised for automated RT-qPCR COVID-19 testing based on a standard 40ft shipping container. Using open-source liquid-handling robots and RNA extraction reagents we demonstrate a reproducible workflow for RT-qPCR COVID-19 testing. With five OT2 liquid handlers, a single CONTAIN unit reaches a maximum daily testing capacity of 2400 tests/day. We validate this workflow for automated RT-qPCR testing, using both synthetic SARS-CoV-2 samples and patient samples from a local NHS hospital. Finally, we discuss the suitability of CONTAIN and its flexibility in a range of diagnostic testing scenarios including high-density urban environments and mobile response units.

**Visual abstract:** 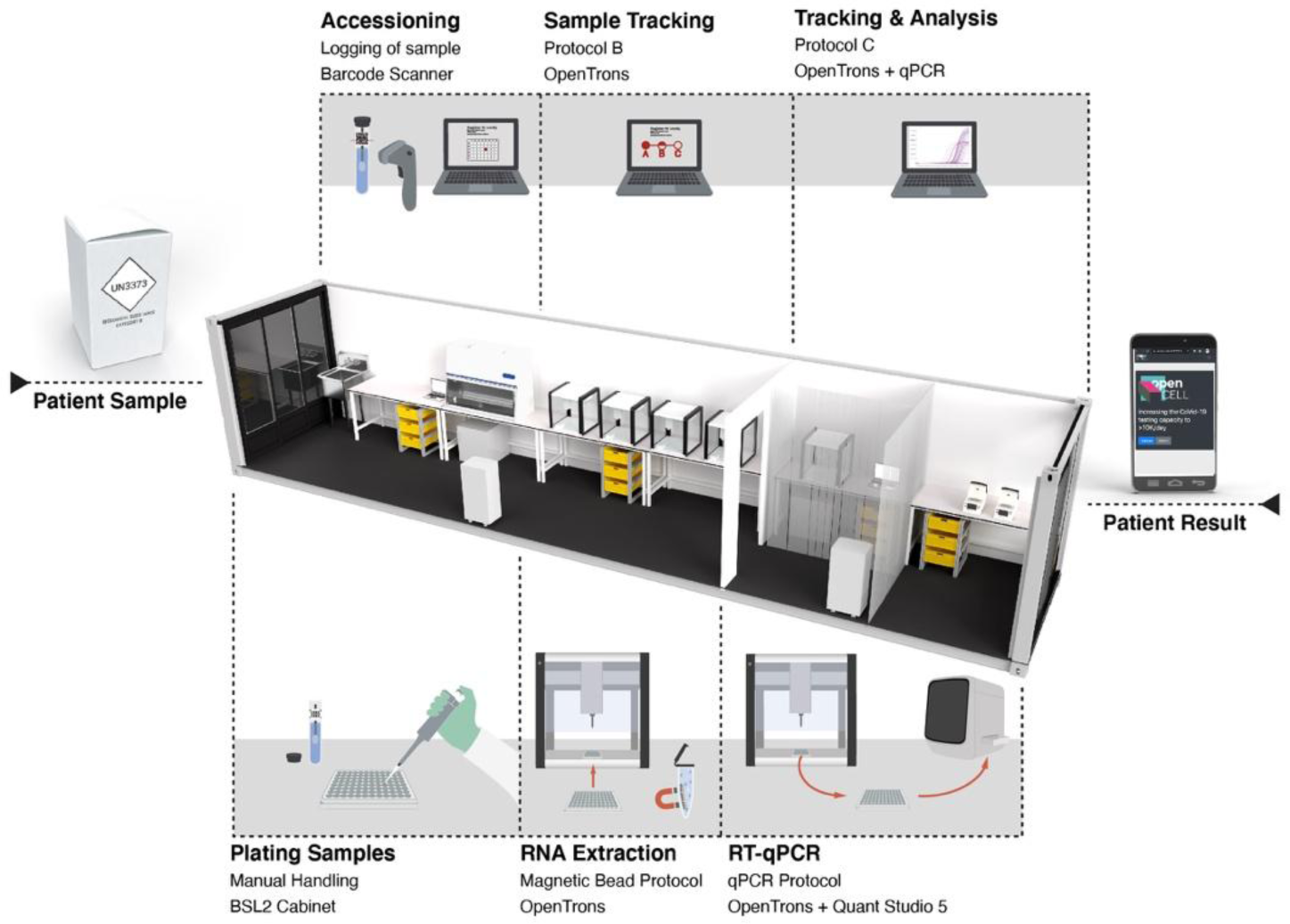

## Introduction

The COVID-19 pandemic presents a diagnostic and logistical challenge. Test, Track and Trace remain the recommended method to control the spread of COVID-19^1^. Nonetheless, the capacity to perform and access diagnostic testing in most countries is still limited. Attempts to scale up testing for COVID-19 have been hampered by limited laboratory infrastructure, supply line disruption and the logistical challenge of population sampling. Since the first reports of the SARS-CoV-2 virus^2^ several studies have proposed new diagnostic approaches that make use of frontier technologies to detect SARS-CoV-2. These include ultra-high-throughput methods based on next-generation sequencing^3,4^, highly informative methods using long-read sequencing^5^, molecular assays that reduce the need for laboratory infrastructure such as loop-based isothermal amplification (LAMP)^6–8^, CRISPR-Cas13 assays^9–11^, and immunology tests that detect the presence of anti-SARS-CoV-2 antibodies^12,13^. These technologies may increase the efficiency and accessibility of SARS-CoV-2 testing in the future but will require regulatory approval. National health organisations still largely rely on reverse transcriptase-qPCR (RT-qPCR)^14–16^.

RT-qPCR is currently the most employed method to test an individual for an active SARS-CoV-2 infection. There have been separate examples of process optimisation for the assay including optimal primer/probe combinations^17^, sensitivity and analytical comparisons of commercially available qPCR kits^18^, and optimised procedures for RNA extractions^19,20^. RT-qPCR technique is sensitive and reliable when performed in a suitable laboratory, however, the rapid deployment of set-ups and workflows capable of coping with the unprecedented volume of testing required has been a challenge for public health organisations internationally. The workflow for preparing a patient sample for RT-qPCR consists of 3 steps: 1) Sample lysis 2) RNA extraction 3) RT-qPCR preparation. In a high-throughput scenario, RNA extraction and RT-qPCR plate preparation are conducted with liquid handling robots, such as the Kingfisher (Thermofisher), CyBio Felix (Analytikjena) Nuclisens EasyMag (Biomerieux). While these robots enable high throughput processing, this approach also requires unique compatible plasticware and proprietary reagents. This has left users exposed to global competition over narrow supply chains and slowed the expansion of testing capability. No report to-date has coupled the complexity of laboratory infrastructure design and the molecular assay process optimisation to propose a standardised solution.

In this work, we present an open-source solution to automated COVID-19 testing - CONTAIN. We have developed automated protocols for RNA extraction and qPCR plate preparation for the open-source and low-cost Opentrons OT2 liquid handlers. The RNA extraction process employs the open-source Bio-On-Magnetic-Bead (BOMB) protocol, which uses DIY reagents and is amenable to automation^21,22^. The use of open-source hardware and reagents allow labs to navigate around limited supply chains^23^. The OT-2 liquid handling robot has a low initial cost relative to other liquid handling robots, can utilise generic plasticware and run protocols written in the coding language Python^24,25^.

We conducted a two-step validation process, first running samples spiked with synthetic SARS-CoV-2 genomic RNA to optimise our process and subsequently patient samples to validate our assay and workflow. We summarise the designs for a portable modular shipping container lab built to a BSL 2+ standard, running up to *2400* tests a day at full capacity. Finally, we discuss three rapid-response scenarios for CONTAIN that reflect a range of needs in this international crisis: 1) a single container to serve a distributed workplace, 2) a single container that can augment the testing capacity of an existing medical facility and 3) a centralised multi-container urban facility.

### A shipping container laboratory

Shipping containers are by definition (1) shippable, allowing a laboratory to be transported to almost anywhere in the world, (2) stackable, allowing laboratory units to be grouped together into large facilities and (3) standardised to increasing reproducibility. The first major challenge in designing a shipping container lab for COVID-19 testing is adapting the container to a biosafety containment level 2+ (BSL 2+) and meeting the ISO15189 standard for a medical laboratory. To achieve a BSL 2+ containment level, it is necessary that the interior surfaces are of hygienic and easily sterilisable materials, and that the features and utilities of the unit make it suitable for the handling of potentially hazardous pathogens (Fig. 1) (for a full list of container alterations see: supplementary materials). The modifications have been completed such that the container can still be shipped as normal ‘freight’ with the original Container Safety Convention (CSC) plates.

**Figure 1:**
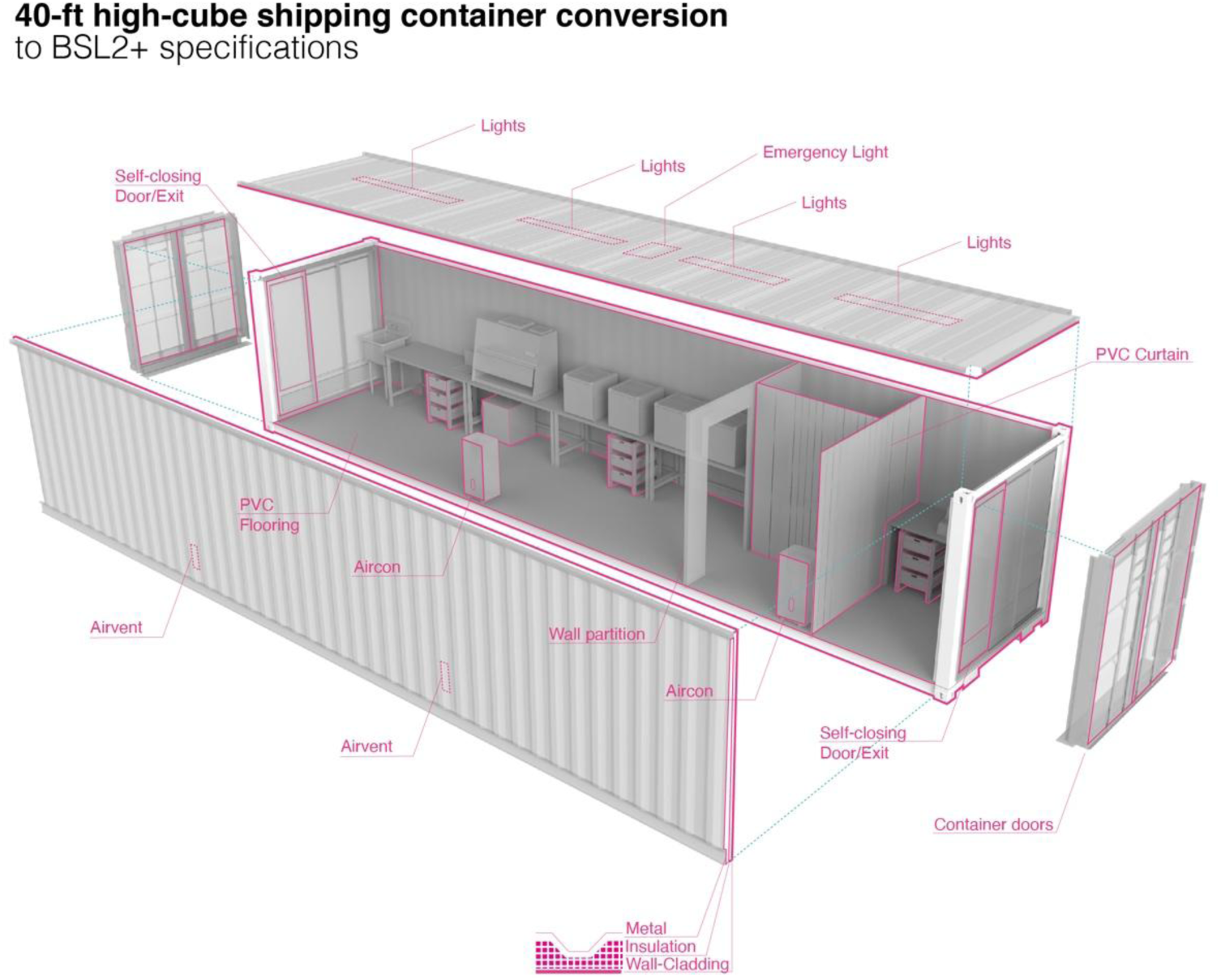
Schematic diagram illustrating the conversion of a standard 40-ft high-cube shipping container into a CONTAIN structure conforming to BSL2+ specifications.

RT-qPCR testing for COVID-19 requires a linear physical layout matching the workflow and cleanliness requirements of the assay. We divided the space into 3 different sections (Fig. 2). Station A: Accessioning, involving the unpackaging and computational logging of samples. Here we provide a microbial safety cabinet, for samples containing active SARS-CoV-2 requiring inactivation. Station B: RNA extraction, outfitted with four OT2s, to conduct multiple simultaneous RNA extractions on samples. Station C: RT-qPCR, containing one OT2 for preparing plates for RT-qPCR, and up to two qPCR machines. Importantly, Station C is not only physically separated from Station A and B but also subdivided further, with a heavy-duty self-sealing PVC curtain insulating the OT2 from the qPCR machines. With this linear physical layout, personnel and samples move from Station A to Station C, changing PPE from B to C, with minimal risk of personal and sample or nucleic acid contamination.

**Figure 2:**
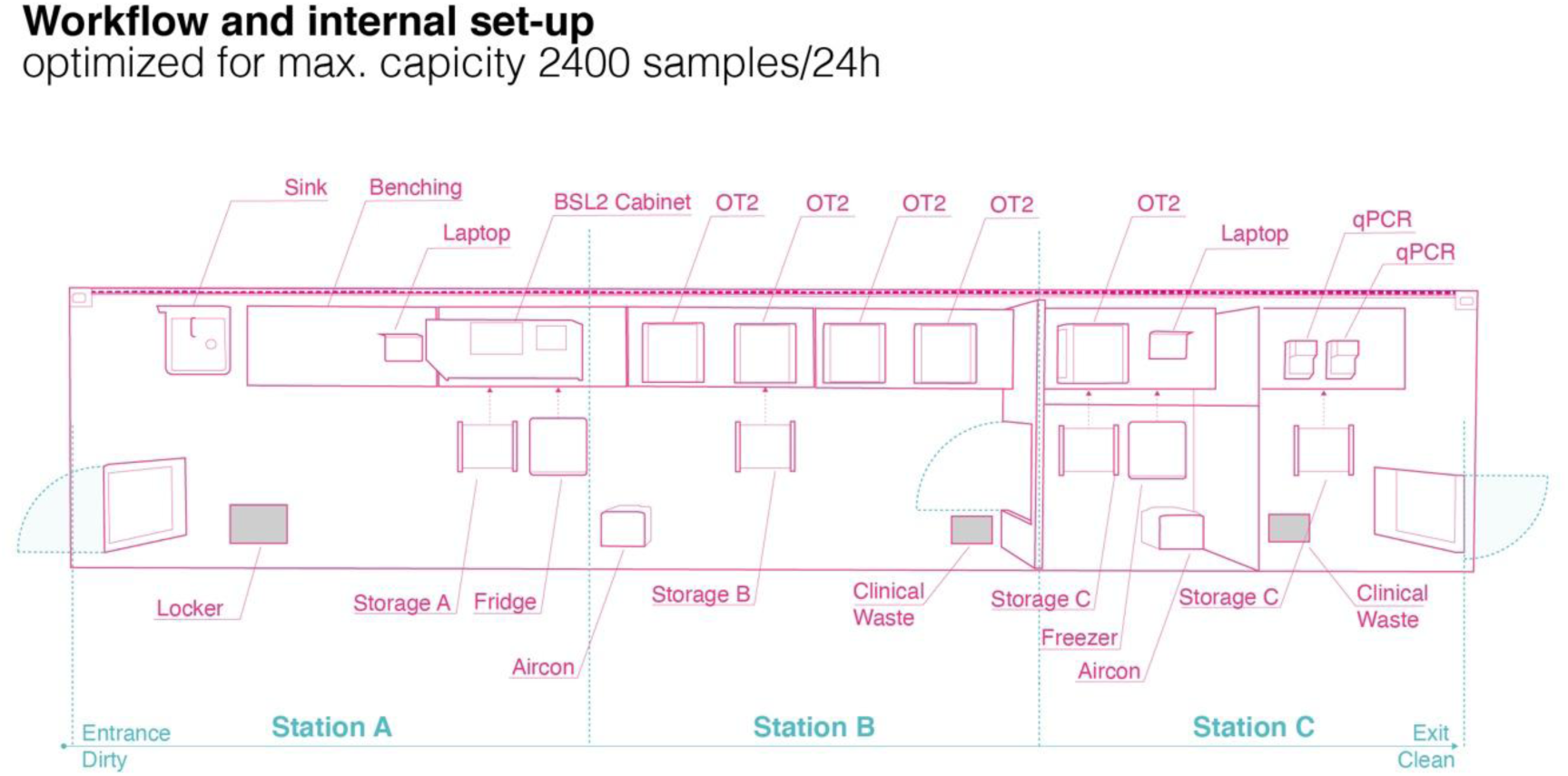
Top-down schematic view of the internal CONTAIN laboratory set up optimized for high-throughput RT-qPCR testing. Workflow is split into three sequential stations, and movement of personnel and labware is strictly unidirectional during operation.

The container lab is designed to plug into the mains and water system of a nearby building or to run from a water tank and generator (Fig. 3). The water system has been designed to prevent contamination of freshwater through an anti-backflow valve and to store wastewater for easy removal by clinical waste disposal services alongside other physical waste. Lab units can be transported by truck to any location and can be placed on a tarmacked space, such as a car park or with minor additional groundwork on uneven ground (Fig. 3). The container lab has been designed to maintain its structural integrity, thereby ensuring compliance with international sea-freight standards. This permits international transport of the lab to any major port (Fig. 3). The structure is also compliant with stacking requirements, meaning multiple containers can be placed securely on top of one another allowing larger structures to be assembled (Fig. 3).

**Figure 3:**
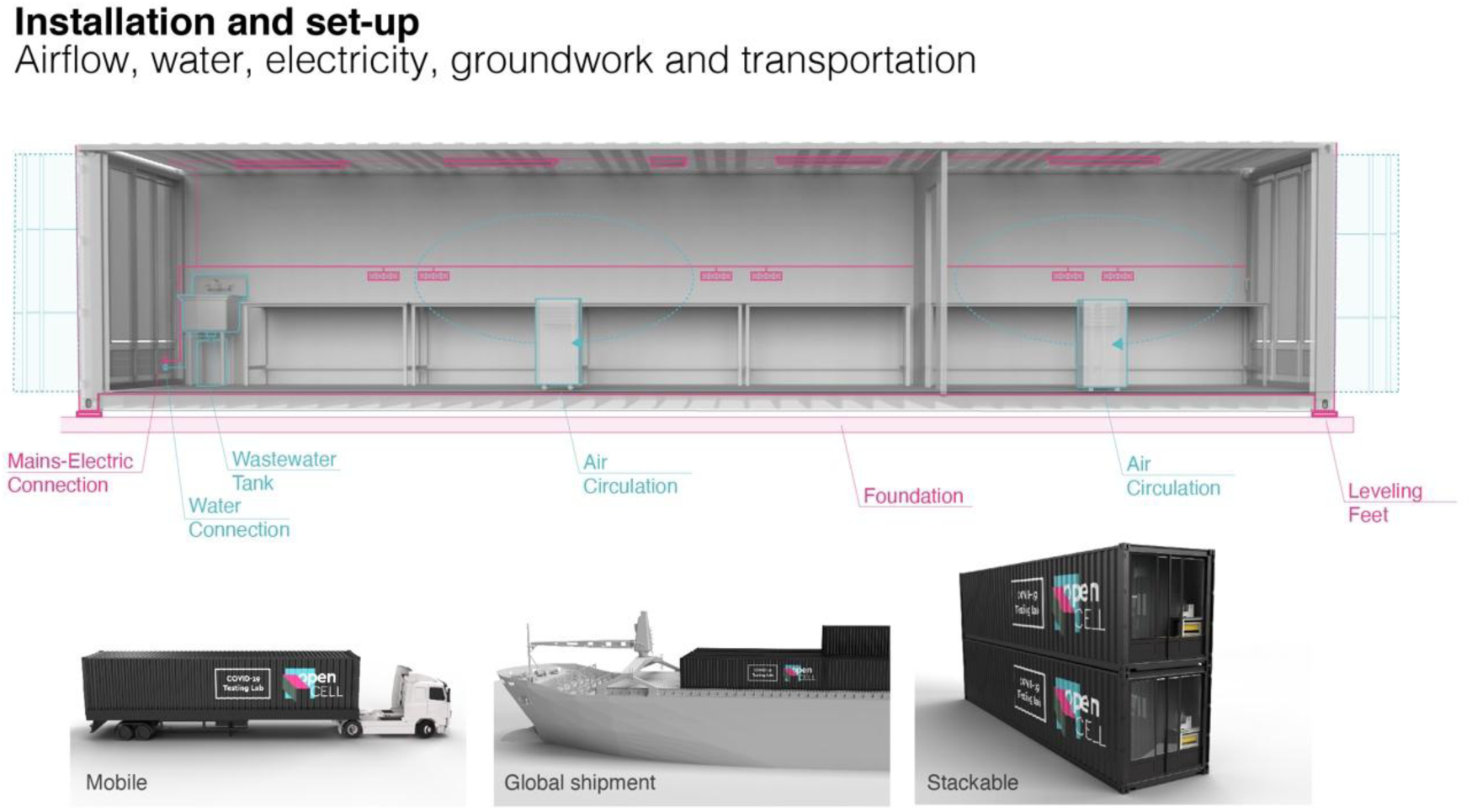
Schematic illustration of CONTAIN utilities and foundation architecture, with examples of transportation and modular stacking of units.

### CONTAIN Automated Assay development

#### Station A

Currently, the most common sampling methods are nasopharyngeal and oropharyngeal swabs. We developed CONTAIN to be a flexible system that works with these, as well as emerging sampling methods, such as saliva^26,27^. After sampling, the swabs can be placed into either viral transport medium or directly into a lysis buffer - which performs lysis of the cellular material whilst inactivating the virus. In Station A, samples are unpacked and logged with a barcode scanner. Then 240 *µ*L of each lysed sample is pipetted into a 2 mL 96 well deep well plate. Once a plate has been filled with 92 samples, the sample plate is passed to Station B.

#### Station B

In Station B, patient samples, already in lysis buffer, are processed to extract RNA for Station C. To develop an RNA extraction method suitable for automation with the OT2, we focused on magnetic beads extraction, using the Opentrons magnetic module attachment. We optimised the BOMB RNA extraction to be amenable to automation with the OT2 (Fig. 4). The automated protocol first isolates the sample RNA on magnetic beads and uses isopropanol and ethanol washes to remove cellular debris and other contaminants from the samples. RNA is subsequently resuspended in nuclease-free water and transferred to a PCR plate for Station C. The automation script is written in Python rather than the Opentrons protocol designer.

**Figure 4.**
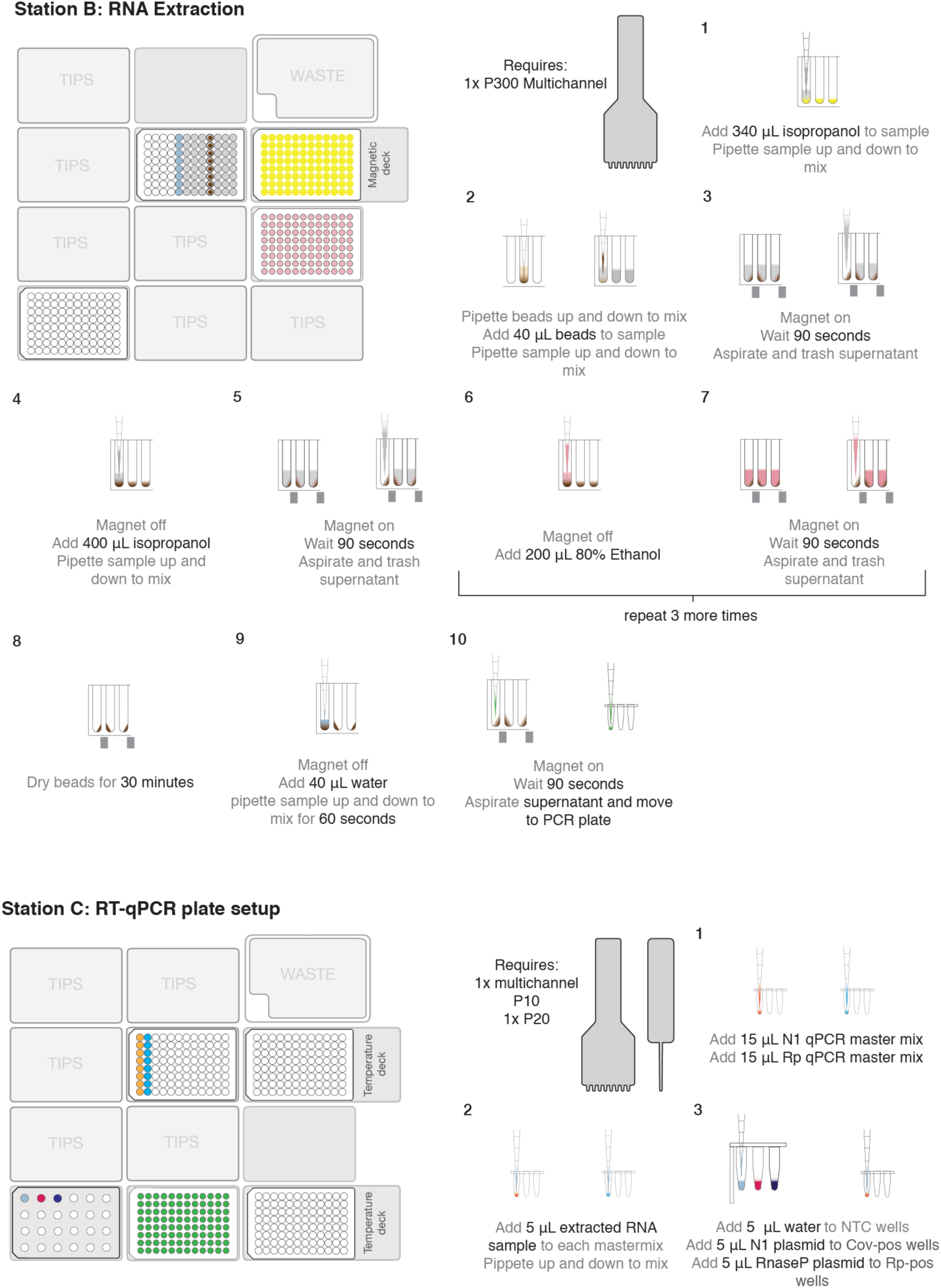
A Graphical breakdown of the automated OT2 protocols. A layout of the OT2 deck for each protocol is shown on the right. Station B is designed to conduct RNA extraction with the BOMB.bio magnetic beads. The protocol makes use of 1 multichannel P300 and uses 200 *µ*L filtered tips to conduct all liquid handling. Colours are yellow for sample in lysis buffer, grey for 100% isopropanol, brown for magnetic beads, pink for 80% ethanol, blue for RNase-free water and green for extracted RNA in RNase-free water. Station C is designed to prepare a plate of extracted RNA for RT-qPCR with a 4x mastermix. The protocol uses both a multichannel P10 and a P20, with 10 *µ*L filter tips. Colours are orange for the N1 master mix, cyan for the RNaseP master mix, magenta for the N1 positive control plasmid, dark blue for the RNaseP positive control plasmid.

#### Station C

In Station C, we have designed an automated protocol for adding one-step RT-qPCR master mixes to the extracted RNA samples and plating each reaction into the final qPCR plate for RT-qPCR (Fig. 4). We based our RT-qPCR reaction around the uniplex SARS-CoV-2 assay from the CDC^14^. We used the N1 and the RNaseP primer-probe mixes in our assay. As we use a uniplex assay, it is necessary to run two RT-qPCR reactions for each sample, one for the N1 target and one for the RNaseP target, which acts as an extraction and sampling control. Both reactions for each sample were conducted on the same plate, however, to conduct RT-qPCR reactions for a full 92 samples, requires setting up two PCR plates for each sample plate. The protocols are open-source and freely available; technical details and protocol code is found on our GitHub (https://github.com/UK-CoVid19/OpentronDev).

### Validation of the CONTAIN automated assay

To validate the automated assay, we required samples with a known amount of SARS-CoV-2 RNA. We therefore developed synthetic samples that could be used to optimise our assay prior to testing with patient samples. These were created by mixing SARS-CoV-2 negative buccal samples held in lysis buffer, with a known amount of synthetic SARS-CoV-2 genomic RNA (Twist Bioscience). As these samples contained human cells, oral secretions and microbes, it is reasonable to assume that RNA degradation occurs as expected rendering these a good test for the reliability of our automated platform.

In the first validation of our automated assay, we used 6 buccal samples, each of which were split into 8 to provide enough human material to generate 46 samples. SARS-CoV-2 genomic RNA was then added to 24 of these samples to a final concentration of 730 copies/*µ*L per sample. These samples were then plated and put through our workflow. The RT-qPCR results showed that for the N1 assay, 21 reactions, out of an expected 24, returned a Cq value of less than 40 and there were no recorded false positives. For the RNaseP extraction control test, 41 reactions returned a Cq value of less than 40, out of an expected 46 (Fig. 5A). As the same amount of RNA had been added to each sample, we estimate the intra-run variation of the CONTAIN workflow (Fig. 5B). Of the 21 samples that returned a positive result for SARS-CoV-2, the mean Cq value returned was 34.1 cycles and the standard deviation of these results was 1.3 cycles.

**Figure 5:**
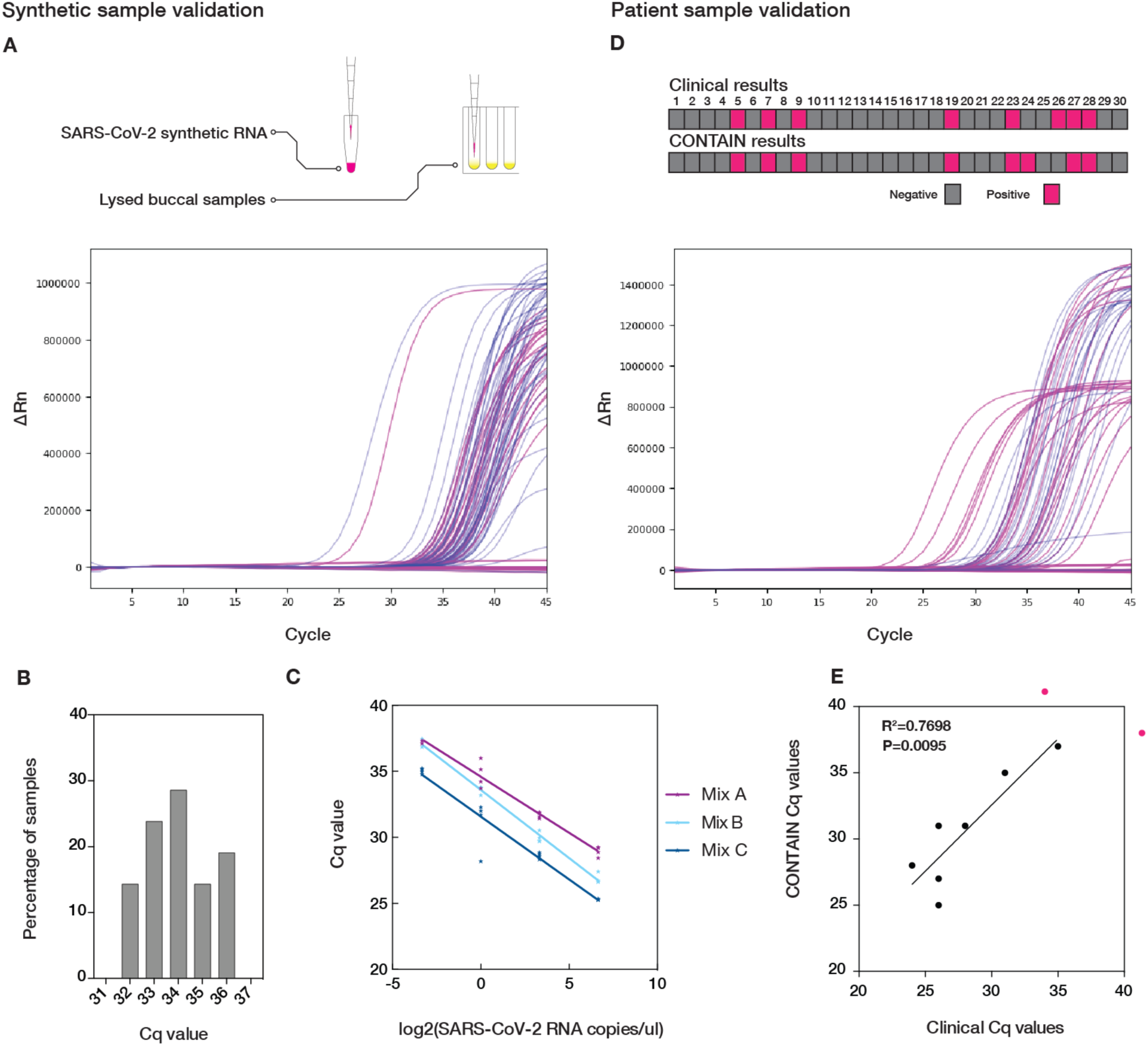
The validation of the CONTAIN automated assay with synthetic and patient samples. A) The amplification curves from the synthetic sample RT-qPCR. The N1 reactions are depicted with pink lines, whilst the RNaseP reactions are depicted with blue lines. B) A histogram of the 21 SARS-CoV-2 positive test results from the synthetic sample tests. C) A comparison of RT-qPCR master mixes using a dilution series of synthetic SARS-CoV-2 genomic RNA. Mix A is the qPCRBIO Probe 1-Step Go (PCR Biosystems), Mix B is the One-Step RT-qPCR mix (Aptogen) and Mix C AgPath-ID™ One-Step (ThermoFisher). D) A comparison of the CONTAIN and clinical results for each of the 30 patients and the amplification curves from the patient sample RT-qPCR. E) The Cq values of the CONTAIN test results compared against the clinical Cq values. The pink dots outside the main graph depicts tests that did not amplify in one of the assays. Only the results that are positive in both the CONTAIN and clinical tests are used in the linear regression.

The initial synthetic validation was conducted using the qPCRBIO Probe 1-Step Go RT-qPCR mastermix from PCR Biosystems. We were also interested in using the N1 primer-probe mix with other one-step RT-qPCR mixes. Using a dilution series of synthetic SARS-CoV-2 genomic RNA, we also tested the AgPath-ID™ One-Step from Thermofisher and the One-Step RT-qPCR mix from Aptogen (Fig. 5C). The test revealed that choice of one-step qPCR mix can impact expected Cq values. Reactions using the AgPath-ID™ One-Step from ThermoFisher had Cq values on average 2.4 cycles earlier than qPCRBIO Probe 1-Step Go from PCR Biosystems.

Following the successful validation of our assay with synthetic samples, we went forward and tested 30 patient samples. The samples were provided to us in lysis buffer, and 8 of the patient samples had already been confirmed by RT-qPCR to be positive for SARS-CoV-2. The samples were run through the CONTAIN workflow, this time using the AgPath-ID™ One-Step RT-qPCR mix from ThermoFisher. The RT-qPCR results showed that for the N1 assay, 8 reactions returned positive, however, one of these had not been detected as positive in the initial clinical testing of the samples, additionally, one sample identified as positive in the initial clinical test but did not amplify in the CONTAIN test (Figure 5D). Finally, we compared the clinical test Cq values with our own CONTAIN Cq values, which revealed that the CONTAIN results correlated strongly (R^2^=0.7698) with the clinical results (Fig. 5E).

### Data Management and Operational Logistics

A digital sample management system has been built to track samples through the testing process, analyse and log test data, and securely communicate test results to the patient and/or healthcare provider.

Samples are tracked with unique IDs linked to a printable QR code (Fig. 6). During sample handling (Station A), sample QR codes are scanned and assigned to a test run with a unique run ID. At Station C, qPCR data is uploaded into the sample management system, and test results (i.e. positive, negative or inhibitory) are automatically generated. Results are communicated using the secure FHIR (Fast Healthcare Interoperability Resources) protocol, enabling data integration with NHS or international healthcare system architectures.

**Figure 6:**
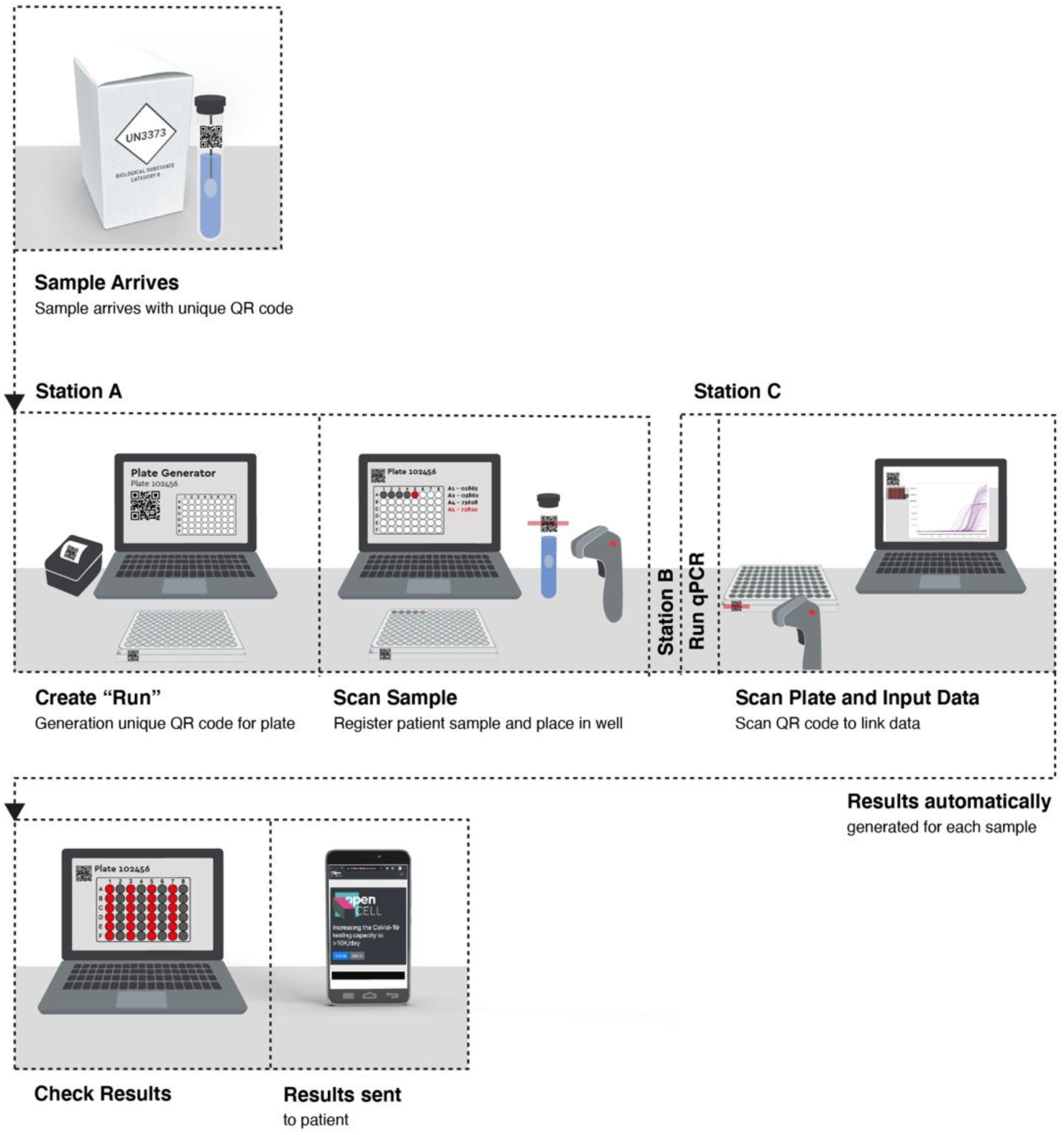
Schematic overview of the sample management system, showing actions performed by laboratory operators.

The system is built using Ruby-on-Rails 5.2 on Ruby 2.4, with a PostgreSQL server to persist data, and operates on a cloud platform. Development is open-source and freely available; technical details and resources are found on our GitHub (ref: https://github.com/UK-CoVid19/opencell-testing) and a more detailed summary can be found in the supplementary materials.

#### Scenarios

The modular and flexible nature of the CONTAIN system would allow it to be used in a variety of conditions. In this section we discuss three possible scenarios highlighting CONTAIN flexibility.

##### University Student and Staff Testing

Universities seeking to resume operation while ensuring the safety of their staff and student population will require regular testing capability. For a large, multi-campus university comprising for example 10,000 staff and 30,000 students, a full-time operation of 10,000 tests per week allows every individual to be tested monthly or more frequently if we consider the majority working from home (Fig. 7). Combined with selective quarantining, this capacity would allow a university to safely resume relatively normal teaching and research practices. Commitment to supporting the health and work of the student body also gives greater peace of mind to the significant overseas student populations who currently face additional challenges in travel, healthcare and accommodation. Two dedicated CONTAIN units can meet a 10,000 test per week capacity, with their mobility enabling them to operate at multiple campuses and accommodation sites throughout the week.

**Figure 7:**
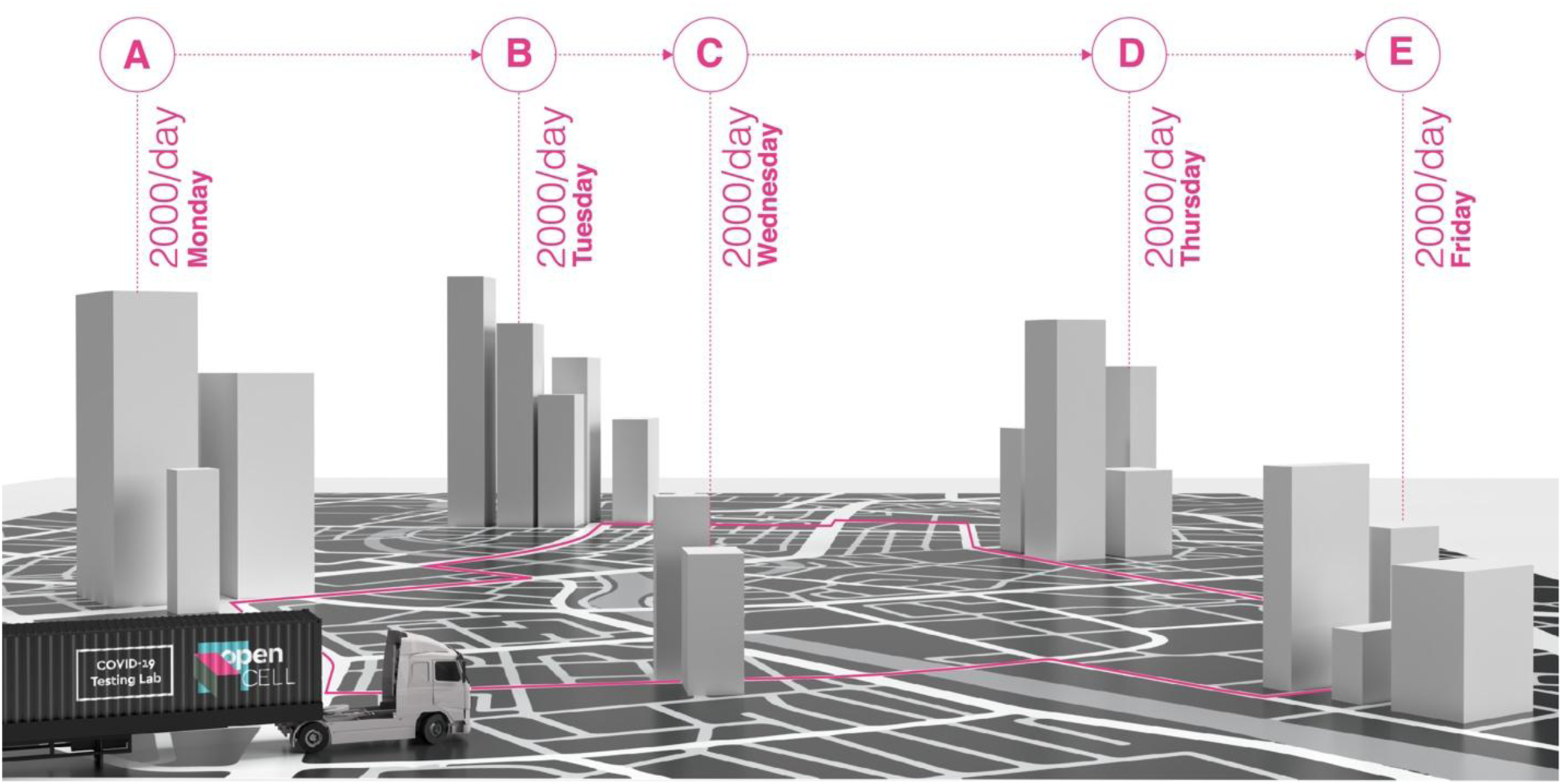
Illustration of a mobile CONTAIN servicing a multi-campus university.

##### Augmenting existing medical infrastructure

Augmentation of existing healthcare infrastructure to meet the surge in demand for SARS-CoV-2 testing requires rapid deployment of suitable (BSL2+) laboratory space. CONTAIN units can be quickly transported to hospital sites and require only a car park area and utilities connection (electricity, water) to be operation-ready. The addition of a dedicated, separate facility enables a hospital to increase the scale of its testing without impacting other essential in-house laboratory work and maintains the safety of the main hospital building (Fig. 8). The flexibility of OpenCell’s CONTAIN system allows hospitals to integrate the unit with their own testing process, existing reagents, waste management system, and/or workforce. CONTAIN units can be transported overseas using standard shipping infrastructure, and additional units may be deployed or moved rapidly between hospitals to react to new outbreaks or surges.

**Figure 8:**
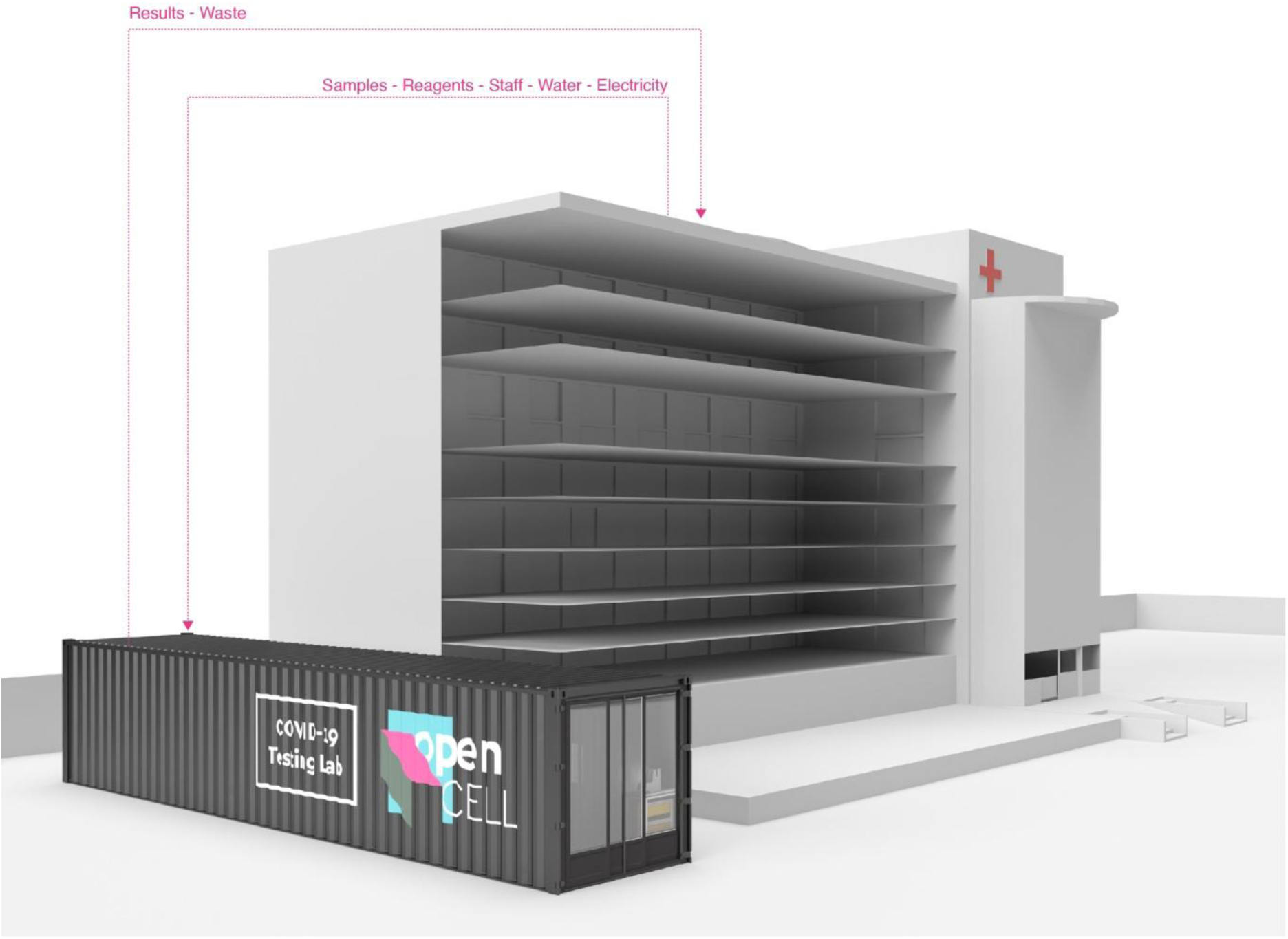
Illustration of a CONTAIN unit augmenting hospital testing capacity.

##### Large Scale implementation: 39 container facility for mass-testing

Large cities make a good logistical case for large centralised testing facilities. In this scenario, we take advantage of the stackability of shipping containers and the modular nature of CONTAIN to envisage a multi-container testing centre [Fig. 9]. The facility, constructed of 39 containers has the potential to run up to 72,000 tests per day at full capacity, and provide office, storage and utility space for the running of the testing centre. This facility could be built rapidly and economically to serve a spike in COVID-19 cases. Once the spike of cases has passed, the facility could then be resourcefully separated back into individual containers and distributed around the country to act as local testing facilities and follow local outbreaks.

**Figure 9:**
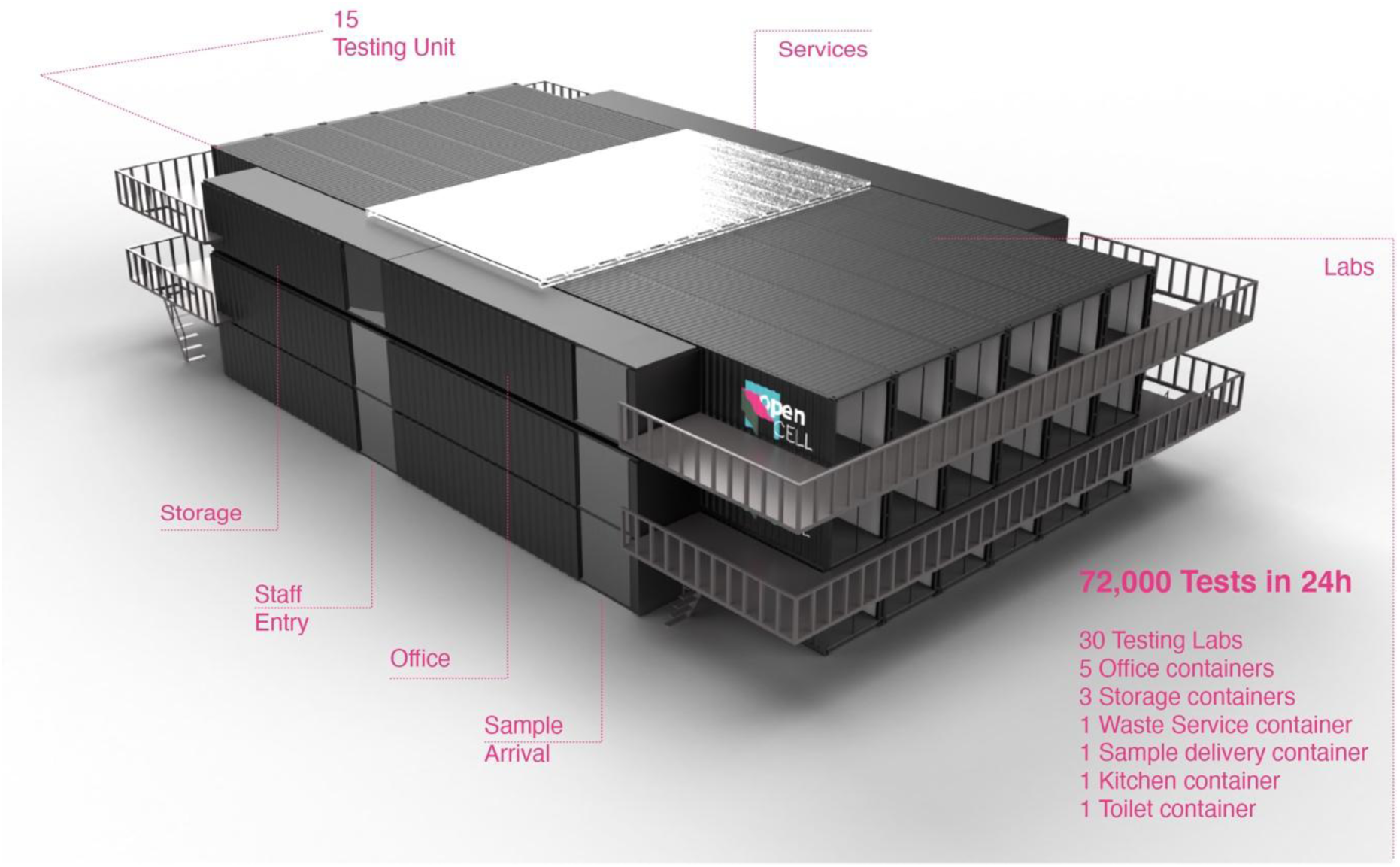
Illustration of a multi-container facility for servicing a large city.

## Discussion

We have demonstrated an open-source shipping container-based laboratory converted to a BSL2+ standard, optimised to perform high-throughput RT-qPCR testing for SARS-CoV-2. Previous work has largely focussed on the implementation of or advances on the testing protocol itself, despite the fact that RT-qPCR is an established and well-characterised protocol, and the challenges faced in the current global crisis in many situations derive from the supporting infrastructure rather than the testing itself. Our laboratories represent a fast, cost-effective solution for the shortage of BSL2+ laboratory space, using standardised and widely available shipping container units. Open-source hardware (e.g. the Opentrons OT2), code, and reagents (e.g. BOMB RNA extraction protocols) permits the use of generic plasticware and easily obtained reagents, circumventing supply chain issues by adapting to local or transient shortages. The shipping container laboratory permits global transport and distribution. Land transport allows laboratories to be rapidly redistributed to locations where they are of the most utility, and little or no preparation of the destination site is required. A custom open-source data management system tracks samples through the system, using the Fast Healthcare Interoperability Resources (FHIR) protocol, an international standard which enables integration into the NHS or healthcare systems in other countries such as the USA. Each laboratory is capable of running up to 2,400 tests/24 hours. In practice, considering for example two 8 hours shift this number is likely to be lower but still over 1500 and compatible with current needs.

The RT-qPCR protocol follows the centre for disease control (CDC) guidelines and is adapted for automation using the OT2 liquid handling robot. As with many components of RNA extraction, magnetic beads kits are in significant demand. We therefore decided to work with the open-source Bio-On-Magnetic-Beads (BOMB) extraction protocol. We made the following major changes to the protocol: removing the DNase digestion step, using a DIY guanidine thiocyanate lysis buffer over TRIzol (Thermofisher) and replacing all mixing steps with repeated pipetting. These changes removed the need for a centrifuge for phase separation and a fume hood for volatile toxic compounds. We also reduced ethanol wash volumes from 400 `μ`L to 200 `μ`L volumes to reduce pipetting steps.

When automating the modified assay, we encountered some issues that required troubleshooting. Due to the precision required at various points of this process, we wrote the OT2 protocol in Python. There is insufficient deck space in the OT2 to house the number of filter tip racks required by the protocol. To circumvent this, a box of 96 filter tips is mapped to the 96 well sample plate, and each tip is reused throughout the isopropanol and ethanol wash steps. This significantly reduces the number of tips required to extract a 96 well plate of samples. We also found that the total removal of ethanol from the final wash step was crucial to allow beads to dry. To achieve this, aspiration of ethanol is performed with the filter tips placed directly at the bottom of the well. Since this sometimes results in the loss of some magnetic beads, the tips are only placed this low for the final wash step. To reduce the chance of cross-contamination, all liquid movements are followed by an additional aspiration of 10 `μ`L of air which acts as a buffer zone. Additionally, the OT2 deck layout has been optimised to minimise the pipette head from crossing over samples. Finally, slightly uneven volumes were occasionally observed when aspirating liquid in the multichannel pipette, we found this to be due to incomplete voiding of liquids. To solve this, excess volume, instead of expected liquid volume, is dispensed.

In developing the automated protocol for Station C, we were aware that incomplete expulsion of liquids from pipettes could be a major source of variation. We therefore followed each expulsion with a blowout to remove any remaining liquid. The CDC assay for COVID-19 initially made use of three probes for SARS-CoV-2, N1, N2 and N3. As studies have shown the N1 primer-probe mix to give the lowest false positive rate compared to N2 and N3, we decided to use only N1 in the CONTAIN automated assay^17^.

Work on the CONTAIN project is ongoing and there is scope for improvement on various aspects. That a sample did not amplify in our testing but did in the clinical testing indicates that this sample may have fallen outside of the limit of detection of our assay and that optimising the efficiency of RNA extraction and sensitivity of RT-qPCR will be an important next step. In the near future we aim to test CONTAIN with a larger batch of patient samples, which will allow us to better ascertain the sensitivity and specificity of our assay. Additionally, benchmarking studies would increase the comparability of the CONTAIN automated system with other diagnostic processes currently in use.

Currently, sampling is performed (in the UK) primarily by nasopharyngeal swabs. However, various studies report high yield and quality of SARS-CoV-2 RNA in saliva, a testing medium better suited to self-testing kits^26^. Validation and/or governmental approval of saliva as a testing medium would simplify the sampling process, reducing the reliance either on trained sampling operators and allowing at-home-testing. Redundancy in the sampling protocol would also circumvent current shortages of the specific swabs required for nasopharyngeal sampling. Nasal swabs and sputum samples have also been the subject of preliminary investigation and also show promise for providing alternative sampling options^28^. Additionally, if standardised, plasticware more amenable to automation is used for sample collection, it may be possible to automate Station A (sample preparation), reducing protocol duration further.

Training of laboratory operators and managers is another focus going forward. Most implementations of the RT-qPCR testing protocol rely on expert operators, while our automated protocol, which outsources many of the technically challenging processes to the OT2 robot, may theoretically be run by a technician with generic laboratory experience. Production of a complete ‘laboratory operation manual’ will allow staff to be sourced and trained locally.

Further investigation of alternative protocols and assays is ongoing, and there is a significant utility in generating guidelines for variations on the standard RT-qPCR process discussed here. The availability of reagents, hardware and wetware varies geographically and temporally. Building a database of alternative RT-qPCR protocols provides an important resource for continuous operation of the laboratories. Recent studies have indicated that the RNA extraction step (Station B) could be skipped and without reducing detectability of SARS-CoV-2 during the qPCR testing step^20^. Efficiencies could be made by switching to a multiplex RT-qPCR, which would save on plasticware, time and reagents^29^. Sample pooling, which involves combining multiple samples in a single well at Station A and testing each sample individually only if the combined sample test positive, would significantly increase the number of tests that can be run by a single CONTAIN unit^30^. This approach, however, is more efficient should the expected number of positive tests be below a certain threshold. Finally, as a liquid handling platform CONTAIN could also be adapted to conduct other types of diagnostic testing, such as ELISAs.

## Acknowledgements

We thank Massimo Majora and the DNA electronics team and Anna Pendrix Russel and Zuzanna Brasko of Sixfold Biosciences for lending equipment for the duration of experiments. We would like to thank Phil Jackson and the MedCity team for support and guidance throughout. We would like to thank Will Canine and colleagues at Opentrons Labworks for their equipment and support. We would like to thank Dr. Catherine Moore for an informative conversation and continued twitter output and Dr Rocio Martinez Nunez for their critical feedback. We would like to acknowledge the support of Charles Walford of Stanhope in advising on infrastructure, groundworks and other practical considerations of development. We acknowledge the support of IDT, Twist Bioscience, ThermoFisher Scientific, MedCity, VWR international, Starlab, Youseq, PCR biosystems, Aptogen for expedited delivery, samples and other support.

## Supplementary information

Code is open-source and present on github: https://github.com/UK-CoVid19/opencell-testing Deployment information is present in the README. Any outstanding questions, please open a new issue at: https://github.com/UK-CoVid19/opencell-testing/issues

Detailed protocols for each station, data, and up to date information on the progress of the project can be viewed at: https://www.opencell.bio/covid-19-testing-laboratories

## SHIPPING CONTAINER SPECIFICATIONS

- 40ft tunnel shipping container (∼300 sqft)
- External structure is modified to sea-freight and land-freight standards (different to other papers/approaches) allowing us to ship the unit across the globe
- Structure is modified to standards so they can be stacked (based on approach to use short ends for entry and exit)
- Containers are designed to be easily plugged into the mains and water system of a nearby building or can be run via water tank and generator
  - Electricity: Commando socket on the outside for the electricity 16amp/connector to connect to mains + Back-up generator
  - Water: 3/4 BSP Threaded Union Connection (Standard washing machine fitting) or Hoselock fitting for simple hose attachment with anti-back flow valve to avoid fresh water contamination and to keep within the regulations of a BSL2+ lab
  - Water Tank for water collection collected by clinical waste service
- The containers are designed for a linear workflow to avoid contamination across stations. From dirt to clean.
  - Entry and exit are placed behind the standard doors of a tunnel container to keep structural properties of the container and enable sea shipment
  - For an optimized linear workflow
  - Doors are self-closing to keep within the regulations of a BSL2+ lab

### Interior fit-out to reach BSL2+

- Closed PVC foam insulation (97% Thermal efficient)
- Hygienic wall-cladding on wall and ceilings
- Hygienic Safety Flooring (chemical resistant and heavy-duty)
- 8 LED strip lights for lighting
- 1 Emergency Light
- 2 Portable BTU 12000 air conditioners with outlets on the site
- Watertank for water collection -> waste service
- Ceiling lights including emergency lighting
- Aircon BTU 12000 for separate areas

### Internal layout optimized for 2400 tests in 24h

3 main areas

- Logging and Plating of samples
- RNA extraction
- qPCR and analysis

### Entry area

- Personal locker and PPE
- First Aid Kit and Eye Washing Kit

### Station A: Logging and Plating of samples

- Sink
- Autoclave
- Manual station / bench
- BSL2+ Cabinet
- PC for logging with OpenCell LIMS and COVID19 software
- Storage unit optimized for reagents and consumables used Station A
- Clinical waste bin

### Station B: RNA extraction

- Benching
- 4 OpenTrons with Magnetic Modules
- Fridge
- Storage unit optimized for reagents and consumables used Station B
- Laptop with OpenCell LIMS and COVID19 software

### Station C: qPCR and analysis

#### Area 1: qPCR preparation

- Benching with PVC curtain for separation to avoid cross-contamination
- 1 OpenTrons with 2x Temperature Module
- Storage unit optimized for reagents and consumables used Station C / area 1
- Laptop with OpenCell LIMS and COVID19 software

Additional PVC curtain separation between area 1 + 2 to avoid RNA cross-contamination

### Area 2

- Benching
- Freezer
- 2 qPCR machines
- Storage unit optimized for reagents and consumables used Station C / area 2

### Exit area

- Clinical Waste bin
- Hand sterilisation at exit

### Transportation and placement of lab unit

Lab unit can be placed on even car parks (temporary) without additional ground preparation and be placed on uneven ground with additional groundwork such as 4 concrete pillars as base

Transportation: Lab units can be transported on moved with a trailer to any location / stays movable

### Added points

- Weekly waste collection - waste service provider

Production time estimate: Production of container and interior 2 working days

Protocol was developed / tested in a minimized set-up (1 qPCR, 2 OpenTrons) - 1000 tests / day

## Supplement: Data Management software CONTAIN system

### Companion software to validate and process viral samples from request to notification

As part of the CONTAIN development, there arose a requirement to generate a consistent data flow of the collection, testing and results of samples. Tests would be requested by a given entity, logged onto a Laboratory Information Management System (LIMS) and monitored through all stations of the laboratory with the results collated and notified to relevant parties.

An open source software application^1^ using Ruby on Rails and the PostgreSQL database, deployed on the Heroku Platform-As-A-Service was developed by volunteers over a number of weeks to serve this requirement.

#### The application was designed to satisfy the following requirements

- Logging the requests for tests from permitted entities (entities defined as members of staff or individuals with access to the platform)
- Logging the collection of samples
- A persistent, valid assignment of samples to a 96 well plate in preparation for laboratory steps A, B and C
- Data export of sample arrangements for feeding into other laboratory hardware
- Receipt and processing of qPCR test results against the validated sample assignment..
- Data access by the least-privilege principle.
- Collation and monitoring of test results
- Continual audit of sample processing; time, step and operator based logging.
- Interoperability with other healthcare data management systems.
- Compatible with commodity hardware (laptop / smartphone)
- Reproducible builds and results.

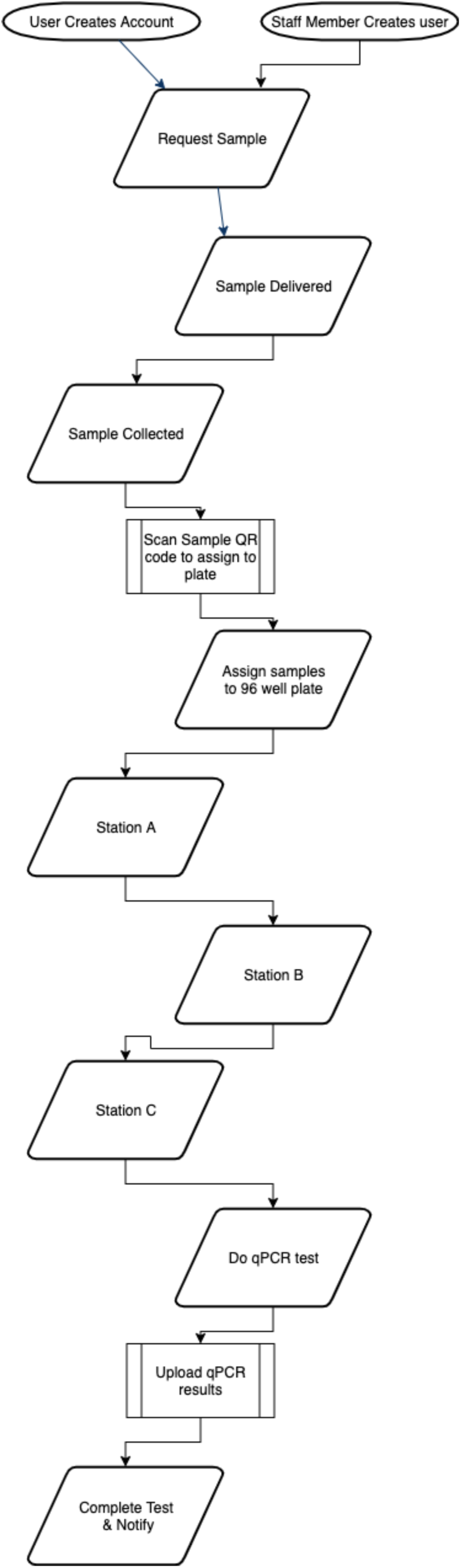

#### Development and Usage

A number of interesting specialized features were developed for this software to suit the above requirements.

Upon the creation of an account, a patient may request a test for themselves or an operator may request a test for a patient and create an account for them.

The application has 3 roles, patient, operator and administrator. A patient can only see their own data, an operator can see limited data and an administrator has access to a special dashboard for making data amendments if necessary.

Once a test is requested, a sample is assigned to the test with a unique identifier. This is encoded as a QR code and is used throughout the process to identify the sample.

The sample has a QR code attached which is then scanned when the samples are assigned to a 96 well plate, the software validates the assignment such that a sample cannot be accidentally assigned to a well twice or in an invalid configuration. A webcam or phone camera can be used to scan the QR code of the sample to ensure simple, consistent assignment. A fallback method with a dropdown list exists when this may not be possible.

The sample proceeds through Station A, B and C upon its plate and is marked on each stage transition. All stages are logged in a history record automatically, based on the user performing the step and the time completed. This can be queried later in the event of a problem.

Upon completion of the qPCR test, the results are uploaded and stored and results assigned back to the original samples and verified.

The results are then interpreted as per the laboratory SOP and an optional feature can be enabled to email patients with their tests results and a custom message.

Data can be exported and downloaded for further collation and dispersion.

Test samples may only monotonically move through the states described in the diagram above, they may also move to a failure state for rework or recollection. The software validates samples can only move in a valid direction.

A test suite verifies the correctness and operation of many of the procedures described above and is included in the code.

Before Station A, data encoding the plate and sample assignment can be exported in CSV or JSON format for input into the Opentrons hardware.

Data collected on patients in this LIMS is as minimal as possible (name, email address).

The decision to use Ruby on Rails for this software was due to the widespread adoption of it in the software development community, mature libraries, ease of development and ease of deployment into many different environments.

#### Integration and extension

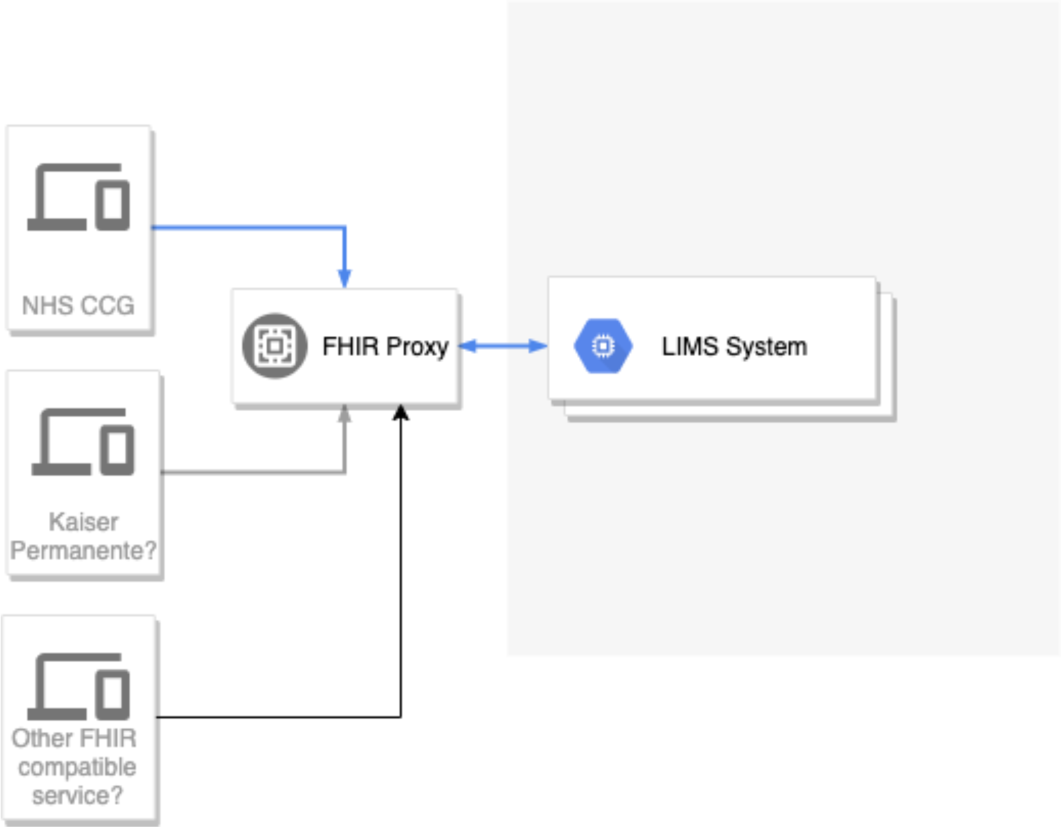

As noted, to interface with public health authorities or wider medical bodies it becomes apparent that a much wider set of data should be collected.

To do this we have investigated the HL7 FHIR protocols which are in widespread use with the NHS^2^ and believe that a logical extension of this software would be to run an FHIR compatible proxy server between the LIMS and outside providers.

^3^

The software already has a comprehensive API which would be able to interact with an FHIR proxy layer.

The application has been left to have a flexible architecture and is open to extension for integration with other modes of requesting tests or dispersion of results.

QR code scanning can be extended to more of the laboratory steps as plates also have unique identifiers.

## Footnotes

1 “UK CoVid-19 · GitHub.” 15 Apr. 2020, https://github.com/UK-CoVid19/opencell-testing. Accessed 19 May. 2020.

2 “National Pathology FHIR Messaging Specifications - NHS “https://developer.nhs.uk/apis/itk3nationalpathology-1-1-0/. Accessed 19 May. 2020.

3 “HL7 UK – HL7 UK Affiliate Site.” https://www.hl7.org.uk/. Accessed 19 May. 2020.

